# A linear neural circuit for light avoidance in *Drosophila* larvae

**DOI:** 10.1101/2021.09.09.459652

**Authors:** Altar Sorkaç, Yiannis A. Savva, Doruk Savaş, Mustafa Talay, Gilad Barnea

**Affiliations:** Department of Neuroscience, Brown University, Providence, RI 02912, USA; Carney Institute for Brain Science, Brown University, Providence, RI 02912, USA

## Abstract

Understanding how neural circuits underlie behaviour is challenging even in the era of the connectome because it requires a combined approach encompassing anatomical and functional analyses. This is exemplified in studying the circuit underlying the light-avoidance behaviour displayed by the larvae of the fruit fly *Drosophila melanogaster*. While this behaviour is robust and the nervous system relatively simple, only bits and pieces of the circuit have been delineated^1^. Indeed, some studies resulted in contradicting conclusions regarding the contributions of various neuronal types to this behaviour^2,3^. Here we devise *trans*-Tango MkII, a new version of the transsynaptic circuit tracing and manipulation tool *trans*-Tango^4^. We implement *trans*-Tango MkII in anatomical tracing and combine it with circuit epistasis analysis. We use neuronal inhibition to test necessity of particular neuronal types for light-avoidance. We complement these experiments by selective neuronal activation to examine sufficiency in rescuing light-avoidance deficiencies exhibited by photoreceptor mutants. Together, our studies reveal a four-order, linear circuit for light-avoidance behaviour connecting the light-detecting photoreceptors with a pair of neuroendocrine cells via two types of clock neurons. Our combined approach could be readily expanded to other larval circuits. Further, this strategy provides the framework for studying more complex nervous systems and behaviours.

## Main Text

Neural circuits underlie all brain functions including processing sensory information and controlling behaviour. Understanding circuit mechanisms necessitates the use of a multi-pronged approach encompassing anatomical and functional analyses. The gold standard in anatomical analysis is EM reconstruction generating a connectome, and much effort has been devoted to producing connectomes of various nervous systems of organisms with increasing complexities. However, even when studying a simple behaviour in a simple organism, analysis of several layers of connected neurons is necessary, a significant challenge when using the connectome data. Further, to truly understand the flow of information in a circuit, one must use functional approaches to manipulate elements within the circuit and observe the consequences. The light-avoidance behaviour, or photophobia, exhibited by larvae of *Drosophila melanogaster* is an example of a robust behaviour in a relatively simple organism. Nevertheless, our knowledge about the neural circuit mediating photophobia is patchy, and at times contradictory^2,3^. To initiate photophobia, light is detected by Rh5 photoreceptors in the larval eye, the Bolwig Organ^2,5^. In the central brain, the prothoracicotropic hormone (PTTH)-expressing neurons are essential for photophobia^6,7^. How these two neuronal types are connected is less clear.

We reasoned that the information regarding light detection by Rh5 photoreceptors could be conveyed to PTTH neurons either directly through synaptic connections or indirectly via other neurons. To reveal whether Rh5 photoreceptors are presynaptic to PTTH neurons, we took advantage of *trans*-Tango, a transsynaptic circuit tracing, monitoring, and manipulation tool^4,8^. While *trans*-Tango has been effectively used to reveal synaptic connections in the adult *Drosophila* nervous system, background noise in larvae^4^ limits its utility in most larval circuits. To solve this problem, we developed a new version termed *trans*-Tango MkII by modifying the ligand construct (Extended Data Fig. 1). Using *trans*-Tango MkII, we found that Rh5 photoreceptors are not presynaptic to PTTH neurons (Extended Data Fig. 2a), indicating the existence of an indirect route.

Which neurons connect the Rh5 photoreceptors to PTTH neurons? The pacemaker clock neurons in the larval visual system are attractive candidates. Expression of the genes *timeless* and *period* (*per*) reveals that the larval visual system comprises nine pacemaker clock neurons: four pigment dispersing factor-expressing lateral neurons (Pdf-LaNs), one Pdf-negative lateral neuron (5^th^-LaN), and two pairs of dorsal neurons: DN1s and DN2s (Fig. 1a)^9^. Inhibition of all clock neurons via expression of the open rectifier truncated potassium channel dORK-ΔC^2^ or of the inward-rectifying potassium channel Kir2.1 (Fig. 1b) results in decreased light avoidance at 1100 or 550 lux, respectively. This observation indicates that at least one of the clock neurons mediates the photophobic behaviour. Indeed, this functional effect is corroborated anatomically as driving *trans*-Tango from the clock neurons reveals direct synaptic input onto PTTH neurons (Fig. 1c).

**Fig. 1:**
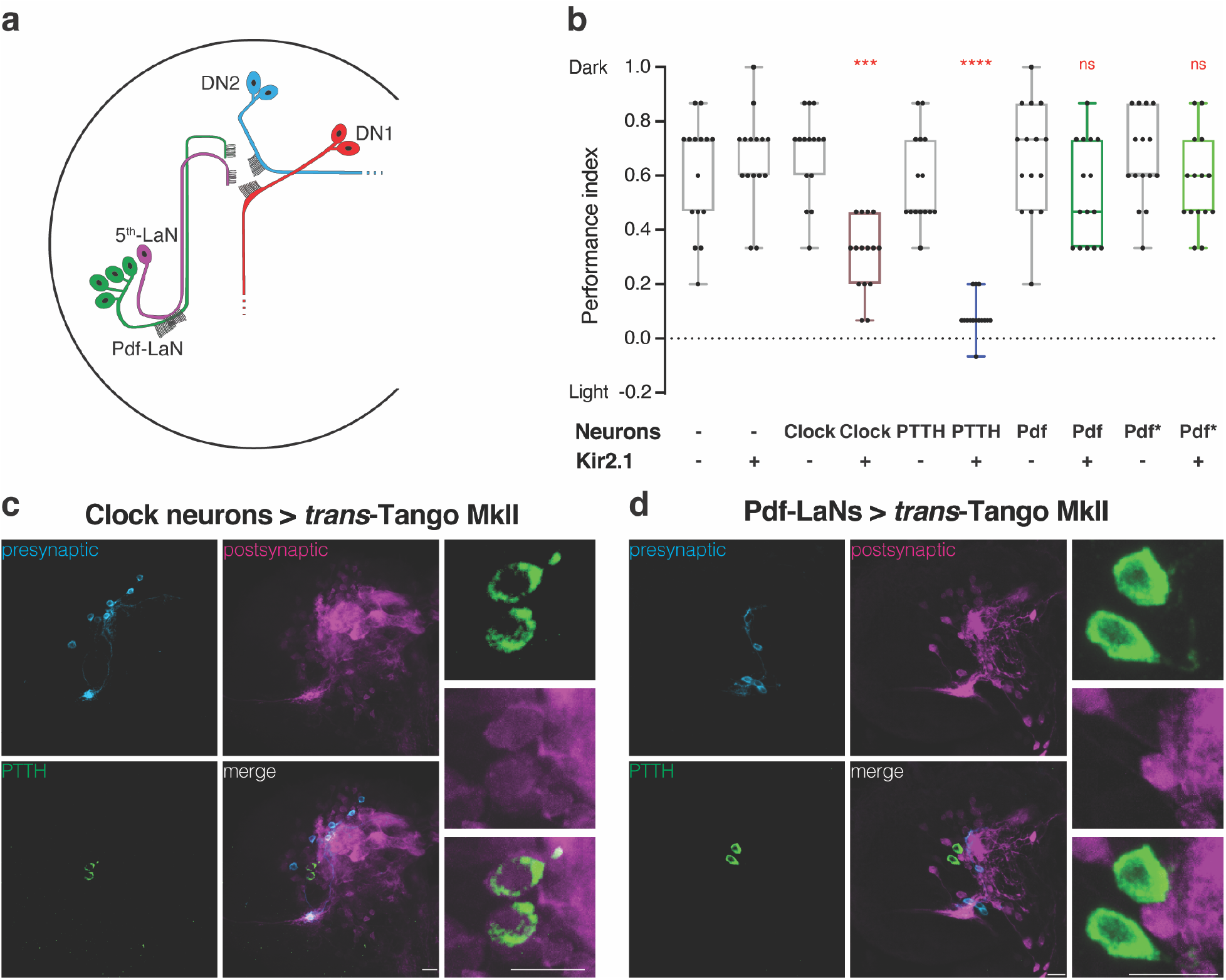
Input from Pdf-negative clock neurons into PTTH neurons mediates light avoidance. **a,** Schematic of clock neurons in the *Drosophila* larval brain. **b,** The effect of Kir2.1-mediated neuronal silencing on light avoidance at 550 lux. Silencing of all clock neurons or PTTH neurons decreases photophobia, silencing of Pdf-LaNs has no effect. Lines represent 75^th^, 50^th^ and 25^th^ percentiles from top to bottom, bars represent maximum and minimum. One-way ANOVA, ns: not significant, ***: p<0.001, ****: p<0.0001. n=15 trials for each group. **c,** Expression of the *trans*-Tango MkII ligand in all clock neurons reveals postsynaptic signal in PTTH neurons. **d,** *trans*-Tango MkII reveals that PTTH neurons are not postsynaptic to Pdf-LaNs. In panels (**c** and **d**), presynaptic GFP (cyan), postsynaptic mtdTomato-HA (magenta), and PTTH (green) are shown. Scale bars, 10μm.

Electron microscopy (EM) reconstruction of the visual system of the first instar larva reveals that Rh5 photoreceptors form synapses with Pdf-LaNs and the 5^th^-LaN^10^. Using *trans*-Tango, we observed that this also holds true in third instar larvae (Extended Data Fig. 2b). Further, Pdf-LaNs have been reported to directly synapse onto PTTH neurons^6^. We therefore sought to silence the Pdf-LaNs by expressing Kir2.1 under the control of two different Pdf-Gal4 drivers. Remarkably, these experiments revealed that Pdf-LaNs are not required for light-avoidance behaviour (Fig. 1b). Therefore, these cells are unlikely the link between Rh5 photoreceptors and PTTH neurons. Indeed, our *trans*-Tango experiments indicate that PTTH neurons are not postsynaptic to Pdf-LaNs (Fig. 1d).

To reveal which of the remaining clock neurons are presynaptic to PTTH neurons, we initiated *trans*-Tango from different subsets. We genetically accessed the 5^th^-LaN with two drivers from the FlyLight collection^11^. These drivers are expressed strongly in the 5^th^-LaN alongside weak and unreliable expression patterns in other neurons^12^. Initiating *trans*-Tango with either driver revealed a faint postsynaptic signal in one of the PTTH neurons, suggesting a potential albeit weak connection with the 5^th^-LaN (Fig. 2a and Extended Data Fig. 3)^12^. We next wished to examine the two pairs of dorsal neurons. However, the drivers used to access DN1s (cry)^13^ or DN2s (Clk9m)^14^ also label Pdf-LaNs. Nevertheless, since Pdf-LaNs do not project onto the PTTH neurons, any postsynaptic signal observed in these neurons would indicate direct synaptic input from DN1s or DN2s. Indeed, initiating *trans*-Tango with either driver reveals that both DN1s and DN2s are presynaptic to PTTH neurons (Fig. 2b, c).

**Fig. 2:**
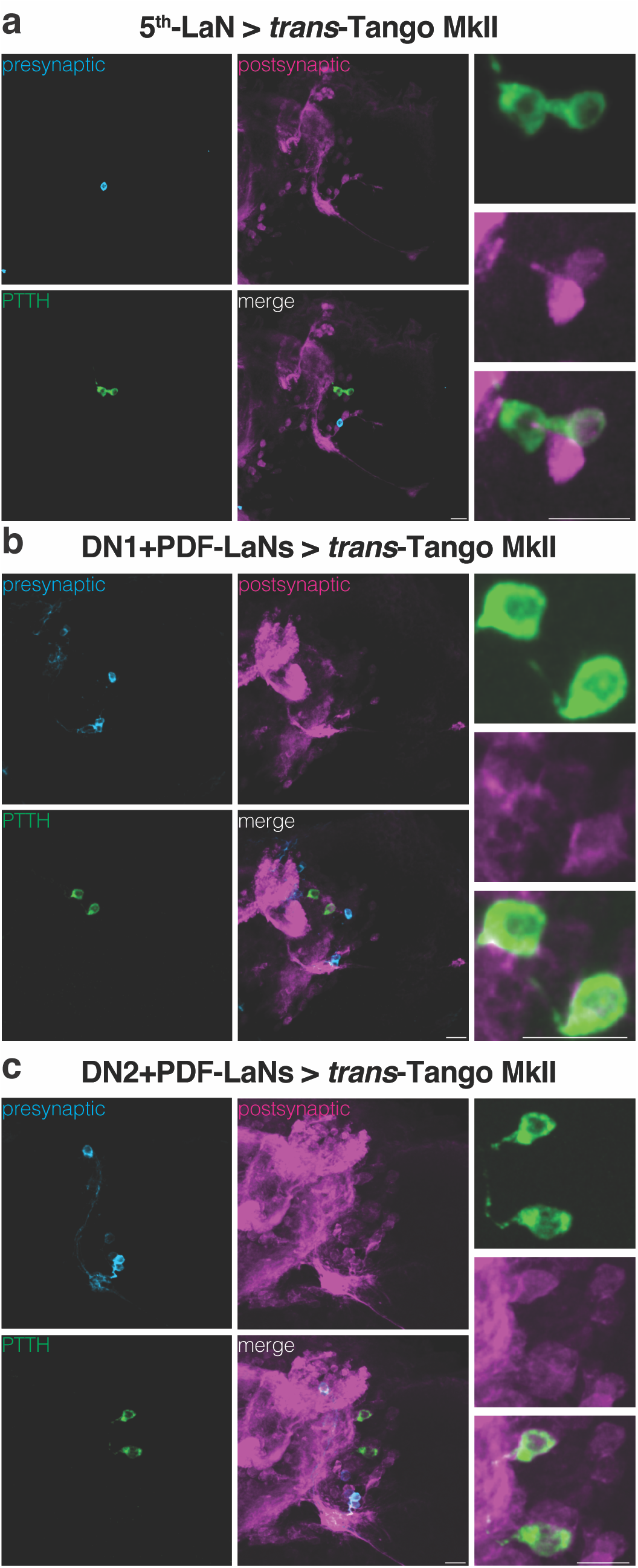
PTTH neurons receive direct input from 5^th^-LaN, DN1 and DN2 clock neurons. **a,** Only one of the PTTH neurons receives input from the 5^th^-LaN. **b, c,** Both PTTH neurons are postsynaptic to DN1s (**b**) and DN2s (**c**). In all panels, presynaptic GFP (cyan), postsynaptic mtdTomato-HA (magenta), and PTTH (green) are shown. Scale bars, 10μm.

We next sought to functionally explore the role of each subset of clock neurons in photophobia using Kir2.1. In accordance with previously published results^3^, we observed that silencing the 5^th^-LaN or DN2s leads to decreased photophobia, suggesting that these neurons are necessary for proper light avoidance. By contrast, silencing of DN1s did not affect light avoidance at 550 lux (Fig. 3a).

**Fig. 3:**
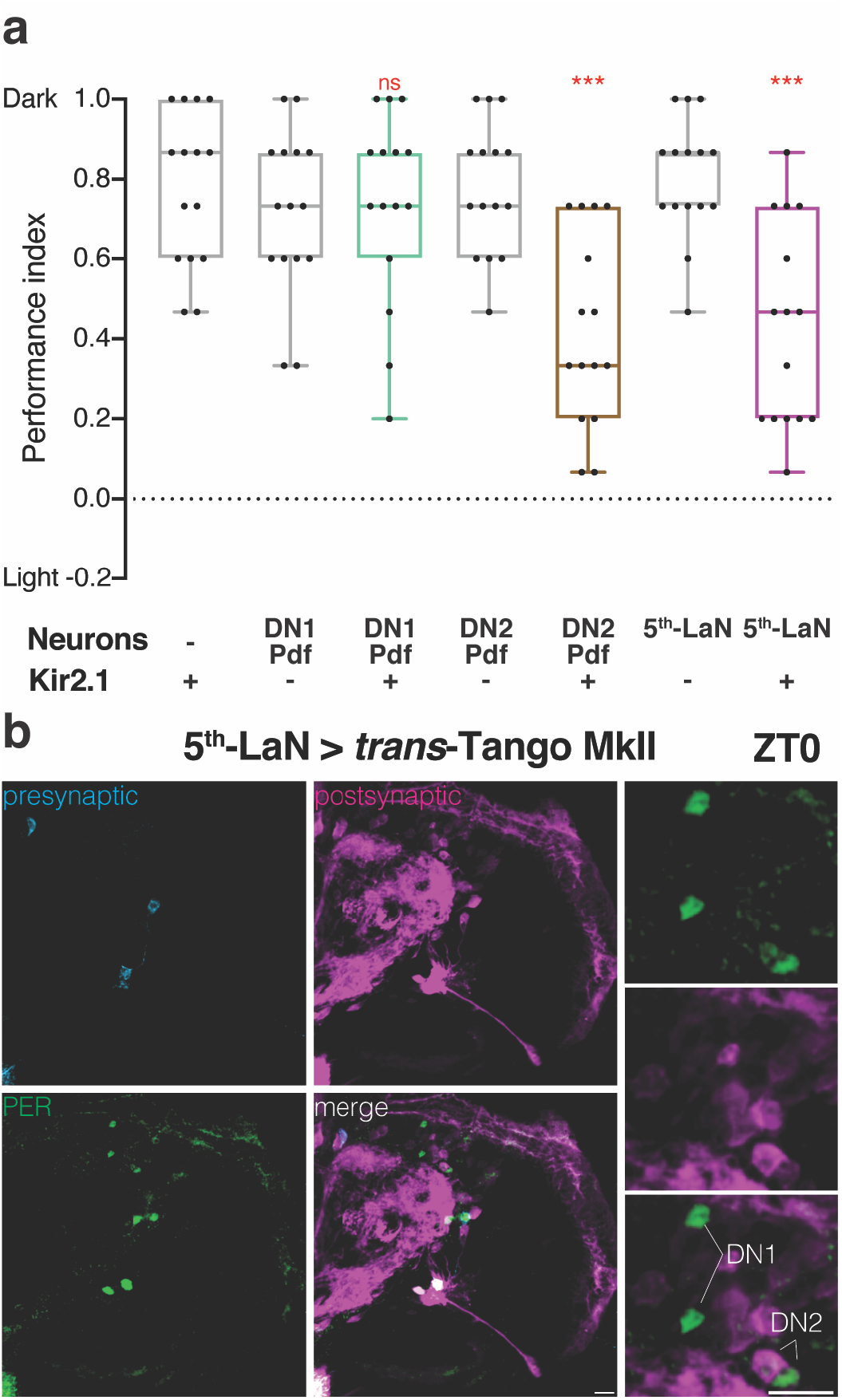
Inhibition of either 5^th^-LaN or its postsynaptic partners DN2s reduces light avoidance. **a,** The effect of Kir2.1-mediated silencing of clock neuron subsets on light avoidance at 550 lux. Silencing of the 5^th^-LaN or DN2s&Pdf-LaNs results in defective photophobia, whereas silencing of DN1s&Pdf-LaNs has no effect. Lines represent 75^th^, 50^th^ and 25^th^ percentiles from top to bottom, bars represent maximum and minimum. One-way ANOVA, ns: not significant, ***: p<0.001. n=15 trials for each group. **b,** DN2s but not DN1s receive direct synaptic input from the 5^th^-LaN as revealed by PER staining in ZT0. Presynaptic GFP (cyan), postsynaptic mtdTomato-HA (magenta), and PER (green). Scale bars, 10μm.

We reasoned that the weak direct connection between the 5^th^-LaN and the PTTH neurons may not be sufficient to convey the light information from Rh5 photoreceptors. Because DN2s are presynaptic to PTTH neurons and are necessary for proper light avoidance, we initiated *trans*-Tango from the 5^th^-LaN to examine whether DN2s constitute an indirect link. We dissected the larvae at zeitgeber time (ZT) 0 when staining with antibodies against PER reveals all clock neurons^9^. We observed that DN2s are postsynaptic to the 5^th^-LaN whereas DN1s are not (Fig. 3b). We confirmed these findings at ZT12 (Extended Data Fig. 4) when PER immunoreactivity is only observed in DN2s^9^. In conclusion, both our *trans*-Tango and neuronal silencing experiments revealed a possible anatomical pathway connecting Rh5 photoreceptors and the PTTH neurons comprising the 5^th^-LaN and DN2s. To further examine this flow of information, we performed functional rescue experiments by activating these subsets of neurons in Rh5 null mutants.

Rh5 mutant larvae are deficient in light avoidance^2,5^. We reasoned that if the 5^th^-LaN and DN2s are indeed downstream of Rh5 photoreceptors, their activation should rescue this deficiency. To test this, we expressed the light-activated cation channel CsChrimson in different clock neurons in the Rh5 null background. CsChrimson can be excited at red wavelengths that are mostly not visible to *Drosophila*^15^. Hence, the red light used to activate CsChrimson would not, itself, cause photophobia. In the functional rescue experiments, we tested the larvae in a modified photophobia assay where half of the plate was dark, and the other half was illuminated with red light to activate CsChrimson. As we anticipated, larvae expressing CsChrimson in the 5^th^-LaN avoided the red-light half of the plate, suggesting that the activation of the 5^th^-LaN is indeed sufficient to induce aversion, and thus, to rescue the light avoidance deficiency of Rh5 null larvae (Fig. 4a). Likewise, we expected that expression of CsChrimson in DN2s would induce aversion since these neurons connect the 5^th^-LaN to PTTH neurons. However, to our surprise, we did not observe aversion to red light when CsChrimson was expressed using our driver for DN2s and Pdf-LaNs (Fig. 4b). Therefore, we decided to examine whether larvae expressing CsChrimson in DN1s would avoid the red-light half of the plate as DN1s are also presynaptic to PTTH neurons. We observed that these animals did not avoid the red-light part of the plate either (Fig. 4c). As is the case for DN2s, our driver for DN1s also expresses in Pdf-LaNs. Hence, we hypothesized that a potential phenotype caused by activation of the Pdf-LaNs might have masked the effects of DN1 or DN2 activation. Indeed, activation of the Pdf-LaNs alone resulted in a slight preference for the red-light half of the plate, rather than avoidance (Fig. 4d), supporting our hypothesis.

**Fig. 4:**
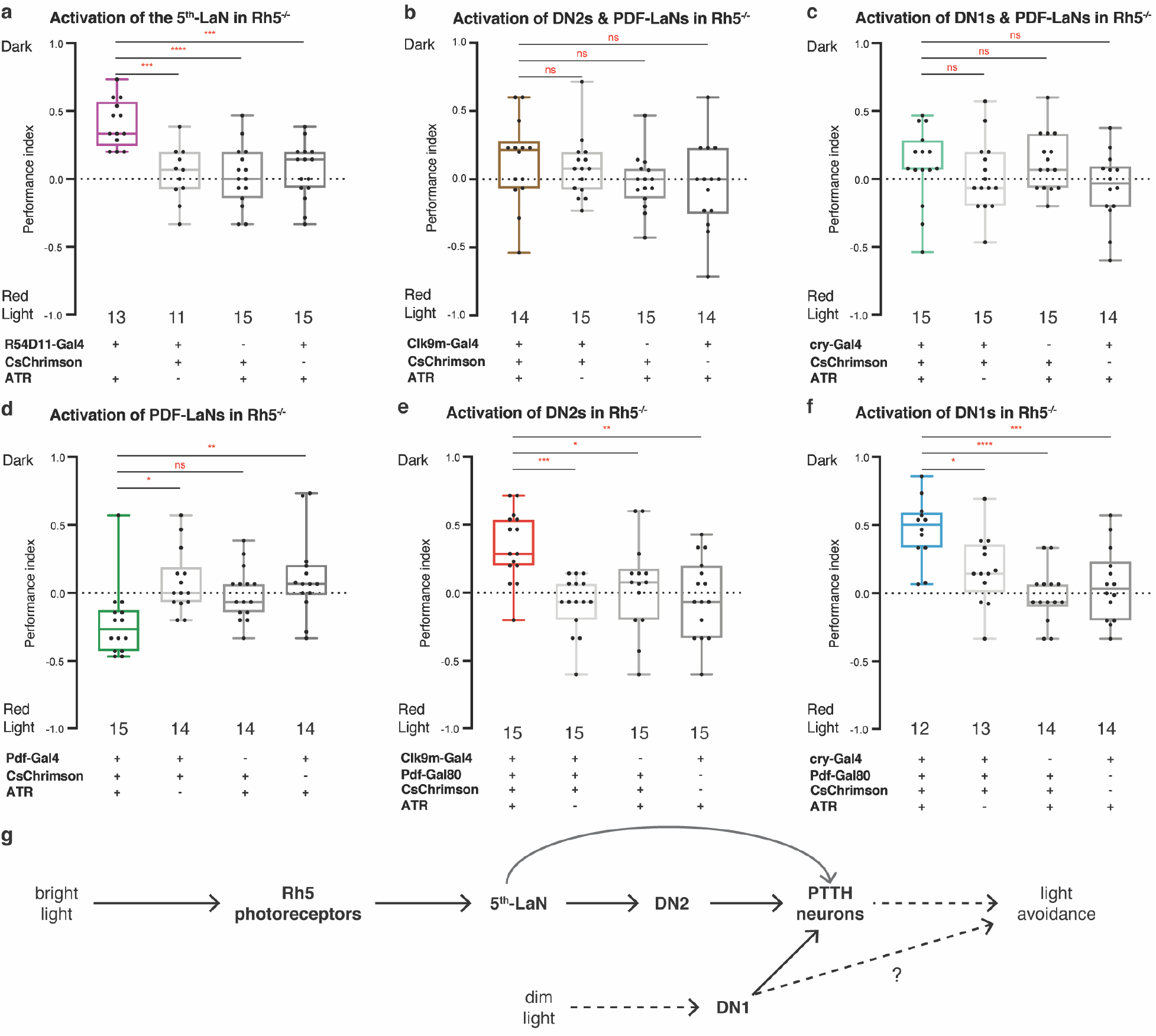
Activation of the 5^th^-LaN, DN2s or DN1s rescues the light avoidance defect exhibited by Rh5 mutant larvae. **a-f** Rescue of the light avoidance defect of Rh5 mutant larvae via CsChrimson mediated activation of specific subsets of clock neurons. Activation of the 5^th^-LaN (**a**), DN2s (**e**) or DN1s (**f**) results in light avoidance. Activation of Pdf-LaNs results in light preference (**d**). No effect is observed when Pdf-LaNs are activated alongside DN2s (**b**) or DN1s (**c**). ATR: all-*trans* retinal. Lines represent 75^th^, 50^th^ and 25^th^ percentiles from top to bottom, bars represent maximum and minimum. One-way ANOVA, ns: not significant, *: p<0.05, **: p<0.01, ***: p<0.001, ****: p<0.0001. The number of trials for each group is indicated below each box. **g,** A schematic showing the flow of information in the neural circuit that mediates the light avoidance behaviour. Bright light activates Rh5 photoreceptors that convey this information to PTTH-neurons via the 5^th^-LaN and DN2s to mediate light avoidance. Dim light indirectly activates DN1s that convey this information to PTTH neurons to mediate light avoidance.

To test the effects of DN2s or DN1s exclusively, we restricted the expression of CsChrimson using the corresponding Gal4 drivers in conjunction with the Gal4-suppressor, Gal80, in Pdf-LaNs (Extended Data Fig. 5). Selective activation of DN2s was sufficient to elicit aversion (Fig. 4e), effectively rescuing the light avoidance deficiency of Rh5 null larvae. These results confirm the neural circuit that connects Rh5 photoreceptors to PTTH neurons through the 5^th^-LaN and DN2s.

Interestingly, selective DN1 activation also led to avoidance of the red-light half of the plate (Fig. 4f). We were puzzled by these results because DN1s are dispensable for photophobia at 550 lux (Fig. 3a) and reduce photophobia at 750 lux^3^. Thus, DN1 activation may cause an ectopic aversion phenotype or affect a different form of photophobic behaviour. Light intensity (*i.e*. dim versus bright light) is an important factor in light aversion^16^. Thus, it is conceivable that DN1s mediate photophobic response to dim light. To test this possibility, we silenced DN1s at 100 lux, and indeed observed that DN1s, but not other clock neurons, are necessary for photophobia at this light intensity (Extended Data Fig. 6). Thus, DN1s may be part of another circuit that mediates avoidance of dim light.

Our study revealed a circuit consisting of four orders of neurons that connect the Rh5 photoreceptors to PTTH neurons via the 5^th^-LaN and DN2s (Fig. 4g). While this circuit mediates the response to bright light, our observation that DN1s are necessary for photophobic response only to low light intensity indicates the existence of an additional pathway for dim light. It is noteworthy that a third, independent system has been reported in which a gustatory receptor mediates photophobic response to high-intensity light^17^. Nonetheless, our results clarify several earlier studies regarding the role of Pdf-LaNs in light avoidance^2,3,5^. In our experiments, Pdf-LaNs are dispensable for light avoidance, yet their activation is attractive. A potential explanation is that Pdf-LaNs may modulate larval photophobia via inhibition^6,7^, especially since adult Pdf-LaNs are glycinergic^18^. In addition, our results contradict a previous study reporting that Pdf-LaNs are presynaptic to PTTH neurons^6^. This study relied on a version of GFP reconstitution across synaptic partners (GRASP) that is in fact not synaptic^19^. Thus, the proposed connection could have been the result of a non-synaptic reconstitution of GFP due to proximity.

Our analysis of the robust light avoidance response in larvae exemplifies the importance of employing a comprehensive approach combining circuit tracing together with neuronal inhibition and activation to test necessity and sufficiency. Our circuit epistasis analysis was made possible by *trans*-Tango MkII, a new version of *trans*-Tango that allows researchers to trace and manipulate neural circuits in *Drosophila* larvae. The combination of a robust and user-friendly genetic tool such as *trans*-Tango with careful functional analysis constitutes a powerful approach that can be readily expanded to studying other circuits and behaviours.

## Methods

### Fly Strains

All fly lines used in this study were maintained at 25°C on standard cornmeal-agar-molasses media in humidity-controlled incubators under 12h light/dark cycle, unless otherwise stated. Fly lines used in this study are in Table 1.

**Table 1:** Fly lines used in this study.

| *Drosophila* Strains | Expression pattern | Associated Figures | Stock |
| --- | --- | --- | --- |
| Clk856-Gal4<sup>23</sup> | All Clock neurons | 1B, 1C |  |
| NP0394-Gal4 | PTTH neurons | 1B | DGRC_K_103604 |
| Pdf-Gal4 | Pdf-LaNs (as Pdf) | 1B, 1D, 4D | Isolated from<br>BDSC_25031 |
| DvPdf-Gal4<sup>24</sup> | Pdf-LaNs (as Pdf*) | 1B |  |
| R54D11-Gal4<sup>11</sup> | 5<sup>th</sup>-LaN, weak and unreliable expression in other neurons | 2A, 3A, 3B, 4A S4, S6 | BDSC_41279 |
| cry-Gal4<sup>13</sup> | DN1s and Pdf-LaNs | 2B, 3A, 4C, 4F, S5A, S6 |  |
| Clk9m-Gal4<sup>9</sup> | DN2s and Pdf-LaNs | 2C, 3A, 4B, 4E, S5B, S6 | BDSC_41810 |
| Rh5-Gal4 | Rh5 photoreceptors | S2A, S2B | BDSC_7458 |
| R19C05-Gal4<sup>11</sup> | 5<sup>th</sup>-LaN, others | S3 | BDSC_48842 |
| Or42a-Gal4<sup>25</sup> | Or42a ORNs | S1A, S1C | BDSC_9960 |
| Pdf-Gal80<sup>26</sup> | N/A | 4E, 4F, S5A, S5B | Isolated from<br>BDSC_80940 |
| w<sup>1118</sup> (5905) | N/A | 1B | BDSC_5905 |
| Rh5[2]<sup>27</sup> | N/A | 4A, 4B, 4C, 4D, 4E,<br>4F, S5A, S5B |  |
| UAS-Kir2.1<sup>28</sup> | N/A | 1B, 3A, S6 | BDSC_6595 |
| UAS-Cshrimson-<br>mVenus | N/A | 4A, 4B, 4C, 4D, 4E,<br>4F, S5A, S5B | BDSC_55136 |
| UAS-myrGFP,<br>QUAS-mtdTomato-HA<sup>29</sup> | N/A | 1C, 1D, 2A, 2B, 2C,<br>3B, S1A, S1C,<br>S2A, S2B, S3, S4 | BDSC_30118 |
| *trans*-Tango<sup>4</sup> | N/A | S1A | BDSC_77123 |
| *trans*-Tango MkII | N/A | 1C, 1D, 2A, 2B, 2C,<br>3B, S1C, S2A,<br>S2B, S3, S4 | This study |

### Generation of Transgenic Fly Lines

The plasmid *trans*-Tango MkII was generated using HiFi DNA Assembly (New England Biolabs #2621) and was incorporated into the attP40 locus using the ΦC31 system as described in the original *trans*-Tango paper^4^. Briefly the hICAM1::dNrxn1 sequences in the *trans*-Tango plasmid were replaced by dNrxn1 sequence amplified from the cDNA clone LP14275 (DGRC #1064347) using the following primers: 5’-atggtaacgggaatactagtCTAGATGGATCGCAAAACTCCTTCTAC-3’ and 5’-ttgttattttaaaaacgattcatggcgcgccTTACACATACCACTCCTTGACGTC-3’. The resulting PCR product was subsequently cloned via HiFi Assembly into the *trans*-Tango plasmid.

### Immunohistochemistry, Imaging, and Image processing

Larval dissections, immunohistochemical experiments, and imaging were performed as described in the original *trans*-Tango paper^4^. Unless otherwise stated, foraging third-instar larvae from vials reared at 25°C were dissected. The antibodies used in this study are: anti-PDF rabbit ^20^ (a gift from Heinrich Dircksen, 1:3000), anti-PTTH guinea pig^21^ (a gift from Michael O’Connor, 1:400), anti-PER mouse^22^ (a gift from James Jepson, 1:50000), anti-GFP rabbit (Thermo Fisher Scientific, A11122; 1:1,000), anti-HA rat (Roche, 11867423001; 1:100), anti-Brp mouse (nc82; DSHB; 1:50), donkey anti-rabbit Alexa Fluor 488 (1:1000), goat anti-rat Alexa Fluor 555 (1:1000), donkey anti-mouse Alexa Fluor 647 (1:1000). Since the *trans*-Tango signal was too weak with R54D11-Gal4 at 25°C, those crosses were set at 18°C for optimization (Fig. 2A, 3B and S4). Resultant images from *trans*-Tango experiments were processed using the Zen software (Zeiss) setting white, black and light corrections in all channels to provide better contrast. In *trans*-Tango figures zoomed-out images represent the maximum projection of the Z-stacks throughout the brains whereas the zoomed-in images were formed using subsets of the Z-stacks for clarity.

### Light Avoidance Behavioural Assay

Larval photophobic behaviour was tested as described previously^2^ with minor modifications. All animals used in this assay have been 6X backcrossed to BDSC_5905. Briefly, foraging early third instar larvae were collected from the food, washed with phosphate buffered saline (PBS) twice and let dry on a surface for 3 minutes. 13 to 16 animals were then transferred along the midline between dark and light halves of a 10 cm round petri dish with 15 mL 1.5% agar solution. Half of the lid was covered with a black tape to form the dark half. The plates were exposed to 100 or 550 lux of white light from above and experiments were run for ten minutes at 25°C. At the end of the ten minutes, larvae on either half of the plate were counted and the preference index was calculated as (# of larvae in the dark)-(# of larvae in the light)/(total # of larvae). For each genotype/condition, at least twelve trials were run over a three-day period. Analysis and determination of significance was performed using One-way ANOVA and subjected to Tukey’s multiple comparison test. Experimental groups were compared to all control groups to determine significance, the lowest pairwise significance is indicated on the figures.

### Optogenetic Rescue Experiments

Optogenetic rescue experiments were run in a similar manner to light avoidance assays with necessary modifications to accommodate for optogenetics. Instead of white light, the light half of the plates were exposed to LED red-light with an intensity of 1600 lux. In addition, parental crosses to obtain experimental animals were set up on standard medium supplemented with 400 μM all-*trans*-retinal (ATR, Sigma #R2500) food or on a standard medium mixed with 100% ethanol for no ATR controls. All animals were kept in 24h dark.

### Data and materials availability

All data are available in the main text or the supplementary materials. Raw data are available upon request. All new fly strains will be deposited to Bloomington *Drosophila* Stock Center.

## Acknowledgments

We would like to thank Alex Fleischmann, Stavros Lomvardas and members of Barnea Lab for critical reading of the manuscript. We are grateful to Heinrich Dirckson, James Jepson, and Michael O’Connor for sharing reagents. This work was supported by National Institutes of Health grants R01MH105368, R21DC014333 R01DC017146 (GB) and Brown University Carney Institute for Brain Science, Suna Kıraç Fund for Brain Science (DS).

## Author contributions

A.S., Y.A.S., D.S. and G.B. conceptualized the story. A.S., Y.A.S., D.S., M.T. and G.B. devised the methodology. A.S., Y.A.S. and D.S. contributed to the investigation and visualization of the results. A.S, Y.A.S., D.S. and G.B. administered the project. A.S., Y.A.S., D.S., M.T. and G.B. wrote the manuscript. The funding was obtained by, and the project was supervised by G.B.

## Additional information

All new fly strains will be deposited to Bloomington *Drosophila* Stock Center. Correspondence and requests for materials should be addressed to G.B. Reprints and permissions information is available at www.nature.com/reprints

