## extended data for "A linear neural circuit for light avoidance in *Drosophila* larvae"

**Extended Data Figures:**

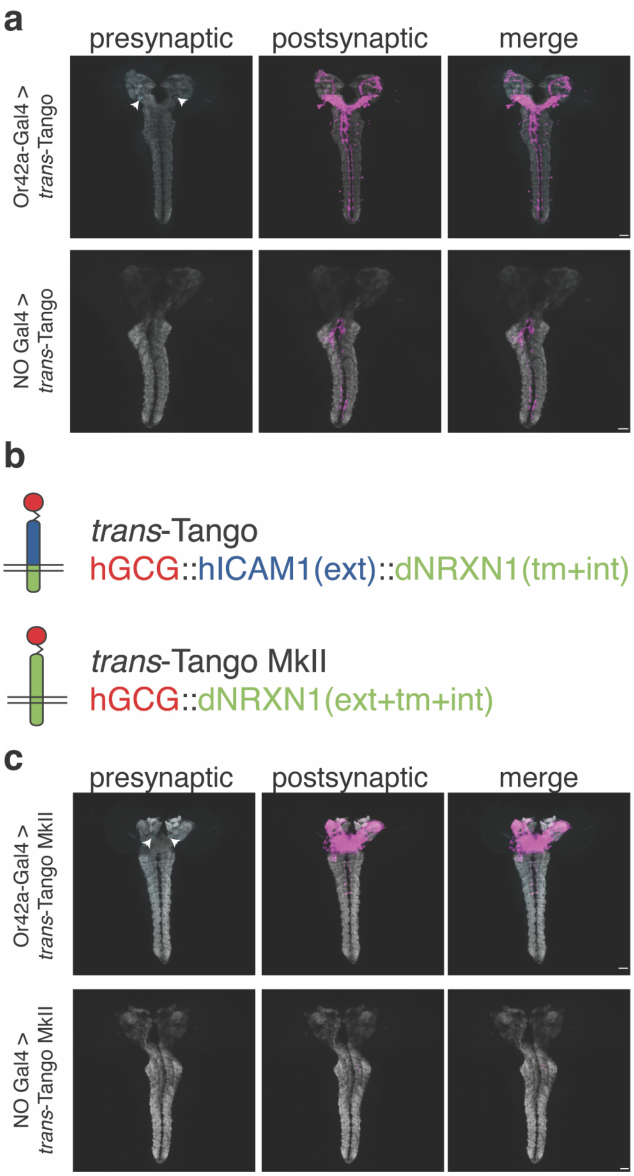


**Extended Data Fig. 1: *trans*-Tango MkII does not have background noise in larval CNS.** **a,** *trans*-Tango experiments using the original ligand results in background noise in the larval VNC in the presence (top) or absence (bottom) of a Gal4 driver. Postsynaptic partners of Or42a-expressing olfactory receptor neurons are revealed. **b,** Comparison of the ligand constructs in *trans*-Tango and *trans*-Tango MkII. In *trans*-Tango MkII the dNRXN1 extracellular domain replaces the hICAM1 extracellular domain that was used in *trans*-Tango[4]. **c,** The background noise in the larval VNC observed in *trans*-Tango is absent in *trans*-Tango MkII with (top) or without (bottom) a Gal4 driver. *trans*-Tango MkII is used to reveal the postsynaptic partners of Or42a neurons. Presynaptic GFP (cyan), postsynaptic HA (magenta), neuropil (grey). Scale bars, 20µm. Arrowheads indicate Or42a glomeruli.

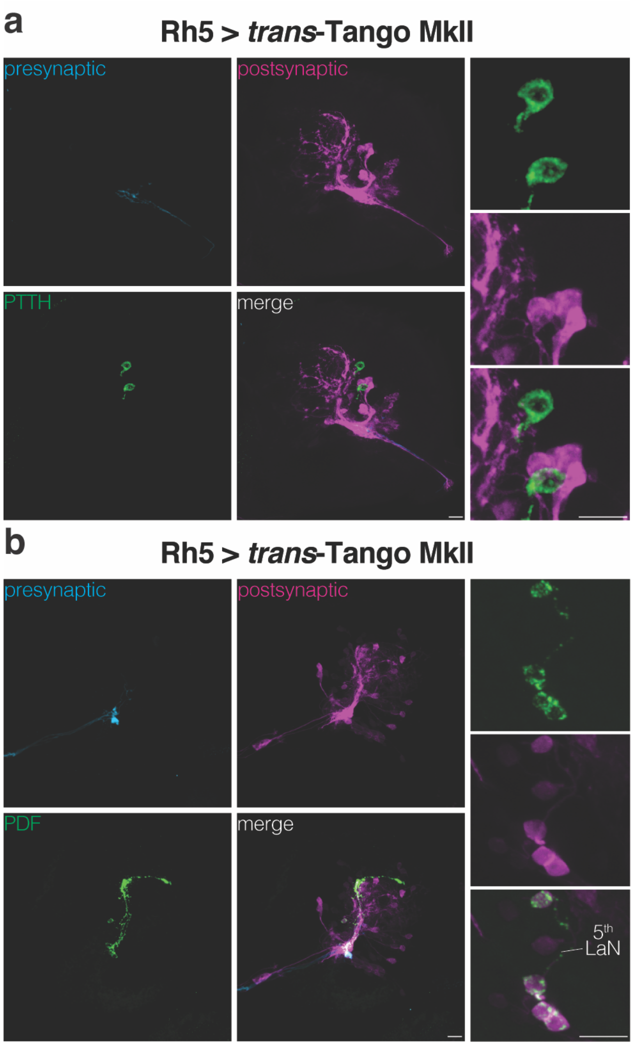


**Extended Data Fig. 2: Rh5 photoreceptors are presynaptic to Pdf-LaNs and the 5$^{th}$-LaN but not to PTTH neurons.**

**a,** PTTH neurons do not receive input from Rh5 photoreceptors. **b,** Rh5 photoreceptors have direct input onto Pdf-LaNs and the 5$^{th}$-LaN. In both panels, presynaptic neurons are revealed by antibody staining against GFP (cyan) and postsynaptic neurons by antibody staining against HA (magenta). Antibodies against PTTH (**a**) and PDF (**b**) were used to reveal the respective neurons (green). Scale bars, 10μm.

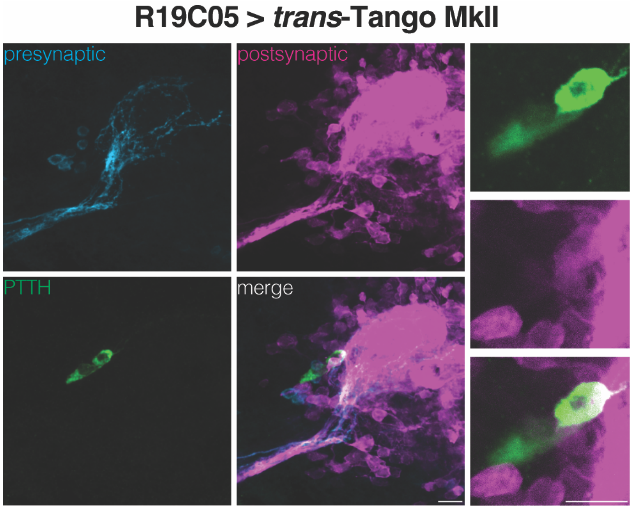


**Extended Data Fig. 3: Only one of the PTTH neurons is postsynaptic to the $5^{th}$-LaN.**

Driving expression of the *trans*-Tango MkII ligand by R19C05-Gal4 confirms that the $5^{th}$-LaN connects to one of the PTTH neurons. Presynaptic neurons are revealed by antibody staining against GFP (cyan) and postsynaptic neurons by antibody staining against HA (magenta). Antibody against PTTH was used to reveal the respective neurons (green). Scale bars, 10µm.

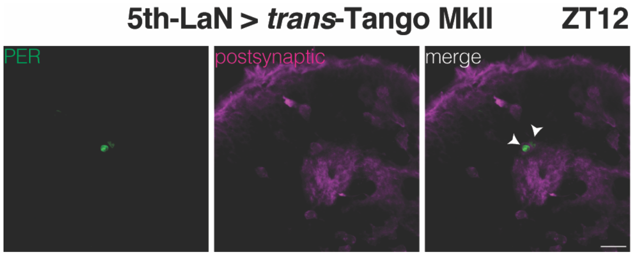


**Extended Data Fig. 4: The 5$^{th}$-LaN is presynaptic to DN2s.**

DN2s receive direct synaptic input from the 5$^{th}$-LaN (R54D11-Gal4) as revealed by PER staining at ZT12. Postsynaptic neurons are revealed by antibody staining against HA (magenta). Antibody against PER was used to reveal the respective neurons (green). Scale bars, 10µm.

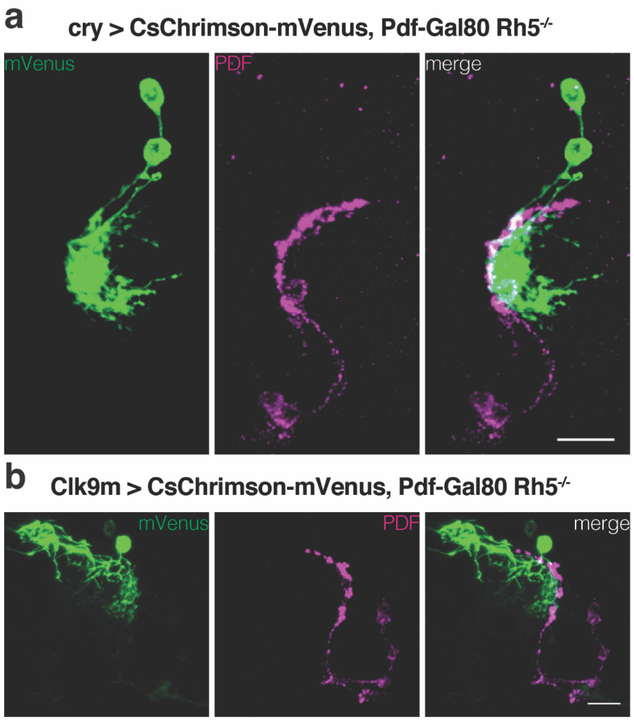


**Extended Data Fig. 5: Selective expression of CsChrimson in DN1s and DN2s in Rh5 mutant larvae.**

**a,** Driving of CsChrimson-mVenus by cry-Gal4 in the presence of Pdf-Gal80 in Rh5 mutant larvae results in selective expression in DN1s. **b,** Driving of CsChrimson-mVenus by Clk9m-Gal4 in the presence of Pdf-Gal80 in Rh5 mutant larvae results in selective expression in DN2s. Antibody staining against mVenus (green) and PDF (magenta) is shown. Scale bars, 10µm.

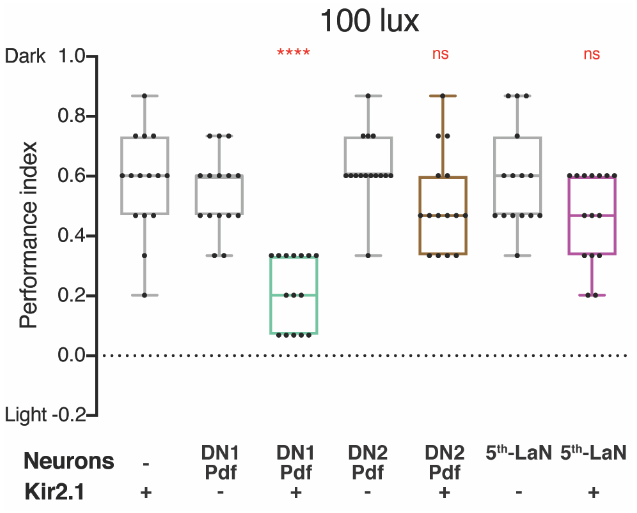


**Extended Data Fig. 6: DN1s mediate light avoidance at 100 lux.**

The effect of Kir2.1-mediated silencing of various clock neuron subsets on light avoidance at 100 lux. Silencing of the DN1s&Pdf-LaNs (cry-Gal4) results in defective photophobia, whereas silencing of $5^{th}$-LaN or DN2s&Pdf-LaNs (Clk9m-Gal4) has no effect. Lines represent $75^{th}$, $50^{th}$ and $25^{th}$ percentiles from top to bottom, bars represent maximum and minimum. One-way ANOVA, ns: not significant, ****: $p<0.0001$. n=15 trials for each group.
